# TAOK2 Drives Opposing Cilia Length Deficits in 16p11.2 Deletion and Duplication Carriers

**DOI:** 10.1101/2024.10.07.617069

**Authors:** Amy Ferreccio, Sujin Byeon, Moira Cornell, Juan Oses-Prieto, Aditi Deshpande, Lauren A Weiss, Alma Burlingame, Smita Yadav

## Abstract

Copy number variation (CNV) in the 16p11.2 (BP4-BP5) genomic locus is strongly associated with autism. Carriers of 16p11.2 deletion and duplication exhibit several common behavioral and social impairments, yet, show opposing brain structural changes and body mass index. To determine cellular mechanisms that might contribute to these opposing phenotypes, we performed quantitative tandem mass tag (TMT) proteomics on human dorsal forebrain neural progenitor cells (NPCs) differentiated from induced pluripotent stem cells (iPSC) derived from 16p11.2 CNV carriers. Differentially phosphorylated proteins between unaffected individuals and 16p11.2 CNV carriers were significantly enriched for centrosomal and cilia proteins. Deletion patient-derived NPCs show increased primary cilium length compared to unaffected individuals, while stunted cilium growth was observed in 16p11.2 duplication NPCs. Through cellular shRNA and overexpression screens in human iPSC derived NPCs, we determined the contribution of genes within the 16p11.2 locus to cilium length. TAOK2, a serine threonine protein kinase, and PPP4C, a protein phosphatase, were found to regulate primary cilia length in a gene dosage-dependent manner. We found TAOK2 was localized at centrosomes and the base of the primary cilium, and NPCs differentiated from TAOK2 knockout iPSCs had longer cilia. In absence of TAOK2, there was increased pericentrin at the basal body, and aberrant accumulation of IFT88 at the ciliary distal tip. Further, pharmacological inhibition of TAO kinase activity led to increased ciliary length, indicating that TAOK2 negatively controls primary cilium length through its catalytic activity. These results implicate aberrant cilia length in the pathophysiology of 16p11.2 CNV, and establish the role of TAOK2 kinase as a regulator of primary cilium length.

## INTRODUCTION

The 593 kb 16p11.2 genomic region between breakpoint 4 to 5 contains 30 annotated genes and is flanked by segmental duplications, making it prone to copy number variation (CNV) such as deletions and duplications^1–3^. 16p11.2 CNVs are associated with autism spectrum disorder^2^, schizophrenia^4^, motor and speech impairment^5^, metabolic body weight disturbances^1,6–8^, and structural abnormalities of the brain^8–10^. Carriers of deletion and duplication in the 16p11.2 locus exhibit certain common developmental, behavioral, and social impairments implying convergent mechanisms caused by reciprocal genetic changes. On the other hand, certain opposing clinical manifestations in 16p11.2 deletion and duplication carriers, such as alteration in brain size and body mass index, present a unique opportunity to dissect function of genes within the locus and determine their individual contributions to 16p11.2 CNV pathophysiology. Carriers of 16p11.2 deletion present with increased brain size or macrocephaly, including larger cortex and thicker corpus callosum; in contrast, duplication is associated with microcephaly, thinner corpus callosum, and smaller cortex^5,9–11^. Another clinical feature is body mass index (BMI). 16p11.2 deletion carriers are prone to obesity (defined as BMI≥30 kg per m^2^) while duplication carriers have low BMI (defined as ≤18.5 kg per m^2^ ^1,6,7,12^. Molecular and cellular mechanisms that could contribute to these gene dosage-dependent phenotypes are unknown, and systematic efforts to map the genetic contributions of these opposing effects are lacking. Even though the syntenic locus in mice offers an excellent genetic model to study this chromosomal disorder, phenotypes of the deletion and duplication mouse are opposing to the human condition^12,13^. However, human induced pluripotent stem cells (hiPSCs) based neuronal and organoid models derived from 16p11.2 deletion and duplication carriers have provided important insights into the pathophysiology of these disorders^14–16^. For instance, excitatory neurons derived from 16p11.2 deletion carriers show hypertrophy, implicating increased neuronal soma and arbor size as contributors to the macrocephaly phenotype^14^.

To identify mechanisms that may contribute to the opposing phenotypes in 16p11.2 CNV carriers, we utilized an unbiased proteomic approach. Our phosphoproteomic analyses of neural progenitor cells (NPCs) differentiated from 16p11.2 deletion, duplication, and control iPSCs revealed differentially phosphorylated proteins enriched in cilia and centrosomal proteins. Primary cilia arise from the centrioles and are specialized sensory organelles present in most cell types including neuronal progenitors, neurons, and astrocytes^17–19^. These organelles are microtubule-based membrane protrusions rich in receptors for signaling molecules that emanate from the cell surface. Defects in ciliogenesis, ciliary length, and trafficking cause several human diseases including neurodevelopmental disorders^20–24^. Intriguingly, signaling through cilia also is a regulator of metabolism, and several ciliopathies are associated with obesity^24,25^. Our study demonstrates that primary cilia are impacted in a gene dosage dependent manner in 16p11.2 CNVs. We show that primary cilium length in 16p11.2 deletion carrier iPSC-derived NPCs are longer than in control NPCs, while duplication carrier derived NPCs have stunted primary cilium. Through knockdown and overexpression screens, we identify the genetic drivers of opposing changes in cilia length in 16p11.2 CNV, and find the protein kinase TAOK2 to be the most significant and dosage dependent regulator of primary cilia length.

TAOK2 kinase is important for neuronal development including dendritic spine development and synaptic function^26–29^. Mutations in TAOK2 are associated with neurodevelopmental disorder such as autism spectrum disorder, and mutations impact both the catalytic and regulatory domains^30^. The canonical isoform of TAOK2 encodes a transmembrane protein kinase localized to the endoplasmic reticulum (ER) and centrosomes, with ER-microtubule tethering property^31^. Despite reports citing its localization at the centrosome^31,32^, the role of TAOK2 in regulation of ciliary length was hitherto unknown. In this study, we find that TAOK2 KO human iPSC-derived NPCs exhibit longer primary cilia, while ectopic overexpression of TAOK2 leads to stunted cilia. Centrosomal accumulation of pericentrin is significantly increased in NPCs lacking TAOK2. Further, balance between anterograde and retrograde ciliary trafficking is altered in TAOK2 KO progenitors as evident from accumulation at the cilia tip of IFT88, a core anterograde ciliary trafficking protein. Finally, using pharmacological inhibition, we show that the catalytic activity of TAOK2 is essential for its role in cilia length regulation. Our study reveals reciprocal alterations of cilia length in neural progenitors derived from 16p11.2 deletion and duplication carriers, elucidates the genetic basis for opposing changes in ciliary length, and reveals a hitherto unknown function of TAOK2 kinase in regulation of primary cilium function.

## RESULTS

### Centrosomal and Ciliary Proteins are Differentially Phosphorylated in 16p11.2 CNV Carrier Derived Neural Progenitors

Three of the 30 genes in the 16p11.2 genetic locus encode for proteins that impact protein phosphorylation: MAPK3 and TAOK2 are serine threonine kinases, and PPP4C is a protein phosphatase (Figure 1A). Therefore, we hypothesized that compared to individuals with 2 copies, the phosphoproteome in duplication and deletion patients would be altered, and that identification of differentially phosphorylated proteins could reveal new insight into the neuropathology of 16p11.2 CNVs. IPSCs from fibroblasts of three 16p11.2 deletion carriers, three 16p11.2 duplication carriers, and three unaffected individuals were obtained (Figure S1A). Differentiation of iPSCs into dorsal forebrain NPCs was induced by dual SMAD inhibition using our previously published protocol (Figure S1B)^14^. NPCs differentiated from 16p11.2 CNV carriers and unaffected control individuals expressed dorsal forebrain progenitor markers PAX6 and NESTIN (Figure S1C). NPCs were grown to confluence, cell lysates were prepared, and equal amounts of protein were utilized for downstream proteomics. Each sample was chemically labelled using the 10plex-Tandem Mass Tag (TMT) reagent, following which samples were combined in equimolar ratio. Phosphopeptides were enriched using TiO2 column, eluted and then analyzed by mass spectrometry^33,34^ (Figure 1B). Mass spectrometry analyses detected 18,894 distinct phosphopeptides from 1927 proteins, of which 43 phosphopeptides from 40 proteins were significantly increased (greater than 2-fold increase) in duplication lines compared to control, and 304 phosphopeptides from 213 proteins were significantly decreased (greater than 2-fold decrease) in the deletion lines compared to control (Figures 1C-D and Table S1). Phosphoproteins downregulated in deletion carriers were enriched in cilia and centrosomal function. Further analyses of downregulated phosphopeptides detected in our mass spectrometry experiment revealed that 13.6% (29 out of 213) proteins were cilia or centrosomal proteins based on their entry in the Cilia and Centrosomal Protein Database^35^ (Figure 1D and Table S2).

**Figure 1.**
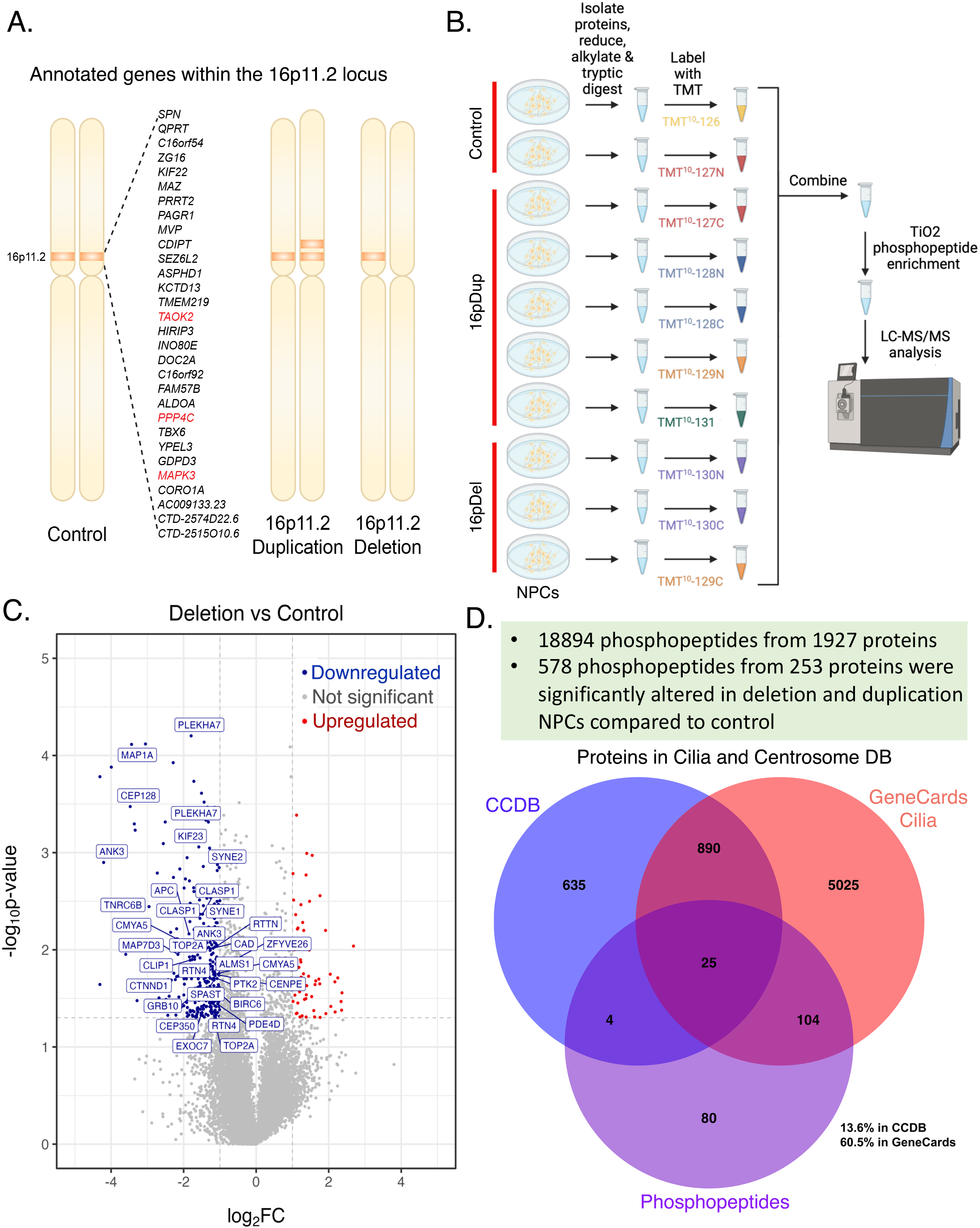
Quantitative Phosphoproteomics of Neural Progenitors from 16p11.2 CNV Carriers Identifies Ciliary Proteins **(A)** Schematic representations of annotated genes within the human 16p11.2 genomic locus that undergo heterozygous deletion or duplication. Genes encoding kinases MAPK3 and TAOK2 and phosphatase PPP4C are highlighted in red. **(B)** Workflow for tandem mass tag (TMT) labeling of protein lysates isolated from dorsal neural progenitors (Pax6, Nestin) differentiated from 16p11.2 deletion and duplication patient as unaffected control individuals. Labelled peptides were enriched for phosphopeptides using TiO2 affinity beads followed by identification through mass spectrometry. **(C)** Volcano plot shows phosphopeptides significantly downregulated (blue) in 16p11.2 deletion samples compared to controls. **(D)** Significantly altered phosphopeptides were enriched in centrosomal and ciliary proteins with 13.6% proteins present in the centrosome and cilia database (CCDB) database and 60.5% proteins had a ciliary function accordingly to GeneCards.

### Primary Cilia in 16p11.2 Deletion and Duplication Patient-Derived Neural Progenitor Cells Exhibit Opposing Defects in Length

To test whether cilia were perturbed in 16p11.2 deletion and duplication carriers, we immunostained NPCs derived from 16p11.2 CNV carriers and unaffected individuals with antibodies against the cilia membrane protein ARL13B and NPC marker PAX6. We found that the primary cilium length in NPCs from deletion patients (mean=3.65 to 4.69µm, n=60 each line) were significantly longer than control (mean=2.36 to 2.55µm, n=60 each line), while cilia length in duplication patient derived NPCs was reduced (mean=1.34 to 1.83µm, n=60 each line) compared to NPCs from unaffected individuals (Figure 2A-B). Cilia length is coupled to the cell cycle^36^. Cilia elongate throughout the G1-S phase of the cell cycle, is disassembled in M phase and then reassembled in G1. Acute serum starvation synchronizes the cell cycle by blocking them in G0 phase where cilia reach maximum length. We found that the reciprocal change in cilia length in NPCs from deletion and duplication carriers persisted, even when NPCs were grown in serum free media for 24hr (Figure 2C). NPCs from both control and 16p11.2 deletion and duplication carriers responded to serum starvation by increasing their ciliary length, however, the differences between these three groups were maintained with increased length in deletion carriers (mean=4.16 to 4.97µm, n>40 each line) and shorter cilia in duplication carriers (mean=2.01 to 2.39µm, n>40 each line) compared to control (mean=2.96 to 3.40µm, n>40 each line). These data together suggest that the deficit in cilia length observed in 16p11.2 CNV carriers are through a cell intrinsic mechanism that is independent of its cell cycle phase.

**Figure 2.**
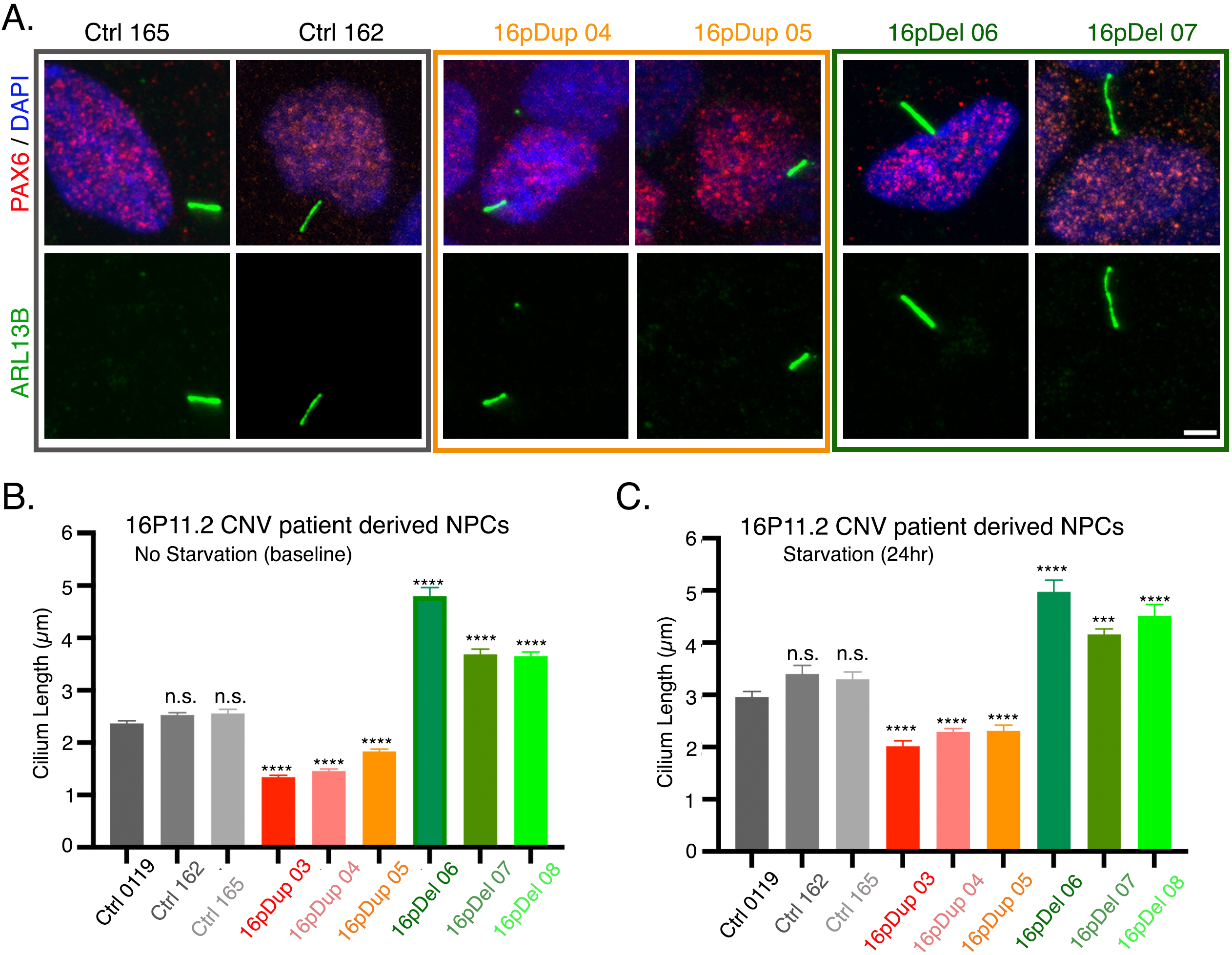
Opposing Cilia Length Alterations in Neural Progenitors Derived from 16p11.2 Deletion and Duplication Carriers **(A)** Representative confocal images of human iPSC derived neural progenitors from unaffected individuals (RMK0162, RMK0165), 16p11.2 deletion carriers (16pDel06, 16pDel07) and duplication carriers (16pDup04, 16pDup05) immunostained for dorsal forebrain progenitor marker PAX6 and primary cilium marker ARL13B. Scale bar is 5μm. **(B)** Length of primary cilia in human iPSC derived neuronal progenitors from unaffected control, and 16p11.2 deletion and duplication carriers, growing in nutrient rich conditions is plotted in microns. **(C)** Length of primary cilia in human iPSC derived neuronal progenitors from unaffected control, and 16p11.2 deletion and duplication carriers, grown in starvation conditions for 24hr before fixation is plotted in microns.

### Cellular Screens Identify Genetic Basis of Altered Cilium Length in the 16p11.2 Locus

To uncover mechanisms underlying altered ciliary length caused by opposing gene dosage changes in the 16p11.2 locus, we performed complementary shRNA and overexpression cellular screens. First, we employed an shRNA screen in the human iPSC lines WTC11 (Coriell Institute) and RMK0119b cells line (UCSF, Weiss Lab) using commercially validated shRNA targeting 23 annotated genes in the 16p11.2 locus. WTC11 or RMK0119b iPSC-derived neuronal progenitors were transfected with GFP-tagged shRNA, starved for 24hr to induce ciliogenesis and then fixed and immunostained with ARL13B antibody 48hr after transfection (Figure 3A). We found that shRNA against TAOK2 (mean=5.43µm, n=26, standard error of mean=0.32) and PPP4C (mean=4.35µm, n= 27, S.E.M.=0.27) significantly increased the cilia length compared to control shRNA (mean=3.43µm, n=29, S.E.M.=0.15) in WTC11 derived NPCs (Figure 3B and 3C). Similarly, in the RMK0119b derived NPCs, shRNA against TAOK2 (mean=4.82µm, n=24, S.E.M.=0.25) and PPP4C (mean=4.29µm, n=16, S.E.M.=0.43) significantly increased the cilia length compared to control shRNA (mean=3.21µm, n=19 S.E.M.=0.19) (Figure 3D). Next, we tested in an overexpression screen, the impact of increased gene expression on cilia length. Briefly, we generated mammalian expression vector with a BFP-tag inserted at the N terminal of TAOK2 and PPP4C along with six other genes in the 16p11.2 locus. NPCs derived from WTC11 line were transfected with either control-BFP construct or constructs expressing the 16p11.2 genes, starved for 24hr, then fixed and immunostained with ARL13B antibody 48hr after transfection to gauge the effect of gene overexpression on cilia length (Figure 3E). Overexpression of TAOK2 had the strongest effect in decreasing cilia length (mean=1.66µm, n=25, S.E.M.=0.08), followed by PPP4C (mean=2.3µm, n=19, S.E.M.=0.15), while control (mean=3.19µm, n=22, S.E.M.=0.16) and none of the other six 16p11.2 genes had a significant effect on cilia length (Figure 3F). These data suggest that gene dosage of TAOK2, and to a lesser extent PPP4C, drive opposing cilia length changes in neural progenitors derived from 16p11.2 CNV carriers.

**Figure 3.**
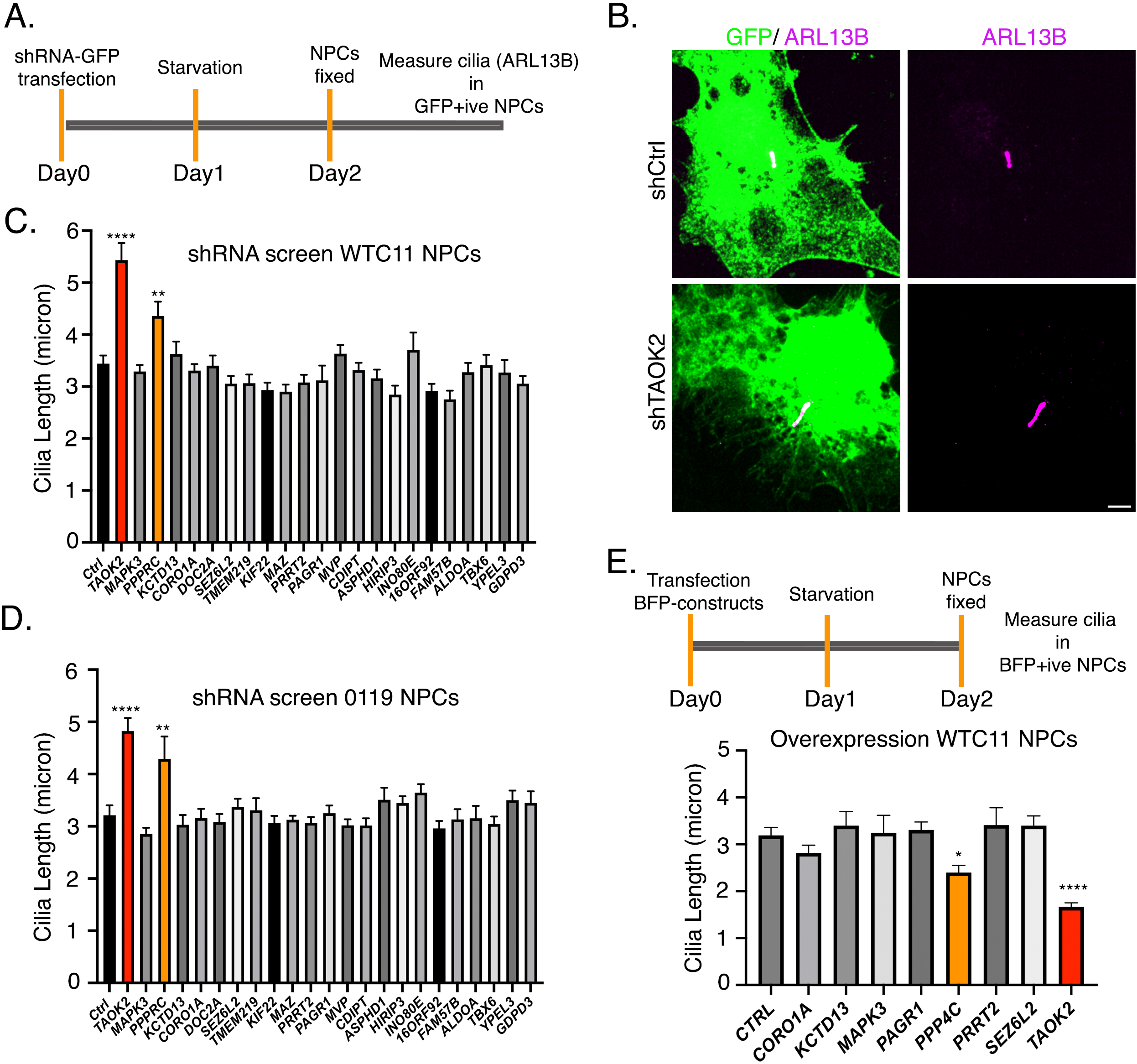
Cellular Screens Identify Genetic Contributors to Altered Ciliary Length in 16p11.2 CNV **(A)** Workflow for shRNA screen involves transfection of WTC11 or RMK0119b iPSC derived neuronal progenitors with shRNA expression constructs (GFP), followed by growth under starvation conditions for 24hr before fixation. NPCs are immunostained for ARL13b to mark the cilium and 60x images are acquired through confocal microscope. **(B)** Representative images of WTC11 derived NPCs transfected with control shRNA and TAOK2-shRNA stained for cilium marker ARL13B (magenta). Scale bar is 2 micron. **(C)** Quantification of cilium length in WTC11 derived NPCs transfected with the indicated shRNA against genes in the 16p11.2 genomic locus. **(D)** Quantification of cilium length in RMK0119b derived NPCs transfected with the indicated shRNA against genes in the 16p11.2 genomic locus. **(E)** Workflow for overexpression screen involves transfection of BFP-tagged expression constructs of indicated proteins in WTC11 iPS derived NPCs, followed by growth under starvation conditions for 24hr before fixation. NPCs are immunostained for ARL13b to mark the cilium and 60x images are acquired through confocal microscope. Quantification of cilium length in WTC11 derived NPCs transfected with BFP-tagged protein expression constructs for the indicated genes in the 16p11.2 genomic locus.

### Cilium Length Deficits in TAOK2 Knockout Neural Progenitors

TAOK2 is a serine threonine kinase localized to the endoplasmic reticulum^31^ that is important for neuronal development^26–28^. In mitotic cells, TAOK2 localizes strongly to the spindle poles^31,32^. To test whether TAOK2 localizes to the cilia and centrosomes in non-dividing neuronal progenitors, we used an antibody against TAOK2 to determine distribution of endogenous in ciliated neural progenitor cells. Immunostaining revealed that TAOK2 was present on the ER membrane in a punctate manner as expected, but was also strongly concentrated at the centrosomes and the base of cilia marked by ARL13B antibody (Figure 4A, and inset). To further validate whether TAOK2, a hit from our cellular screens regulates cilium length, we generated TAOK2 knockout iPS cell lines. We used the AICS0032 stem cell line (Allen Institute), where the centrosomal protein Centrin-2 (CETN2) is tagged with RFP at its endogenous locus, allowing exquisite visualization of centrosomes in stem cell and derivative cells (Figure S3A and S3B). AICS0032 iPSCs were genetically edited using Cas9 and guide RNA targeting exon4 to generate TAOK2 KO cell lines (Figure S3C). TAOK2 KO clones 4.4 and 4.9 were confirmed by sequencing, and absence of TAOK2 protein was validated by western blot (Figure S3D and S3E). Unedited wild type (WT) AICS0032 iPSCs were used as a control. All three lines were karyotypically normal (Figure S3F), and edited iPSCs retained pluripotency as evident by markers NANOG and OCT4 (Figure S3G). Dorsal forebrain neural progenitors were differentiated from WT, TAOK2 KO4.4 and KO4.9 iPSCs using dual SMAD inhibition using our published protocol^14^. TAOK2 KO and WT control NPCs expressed dorsal neuronal progenitor markers PAX6 and NESTIN (Figure 4B) as well as loss of TAOK2 protein in NPCs was confirmed by western blot (Figure 4C). Dorsal neural progenitors from control and TAOK2 KO iPSCs self-assembled into rosettes, and their primary cilia were polarized and projected towards the apical side of the rosette (Figure 4D), mimicking the projection of cilia *in vivo* into the ventricular space. To test whether primary cilium length is altered in TAOK2 KO NPCs, we immunostained NPCs differentiated from WT control and TAOK2 KO4.4 and 4.9 clones with ARL13B to visualize (Figure 4D). We found a significant increase in cilia length in TAOK2 KO 4.4 NPCs (mean=3.52µm, n=151, S.E.M.=0.08), TAOK2 KO 4.9 NPCs (mean=3.35µm, n=170, S.E.M.=0.06), compared to WT control (mean=2.14µm, n=260, S.E.M.=0.04) NPCs (Figure 4E).

**Figure 4.**
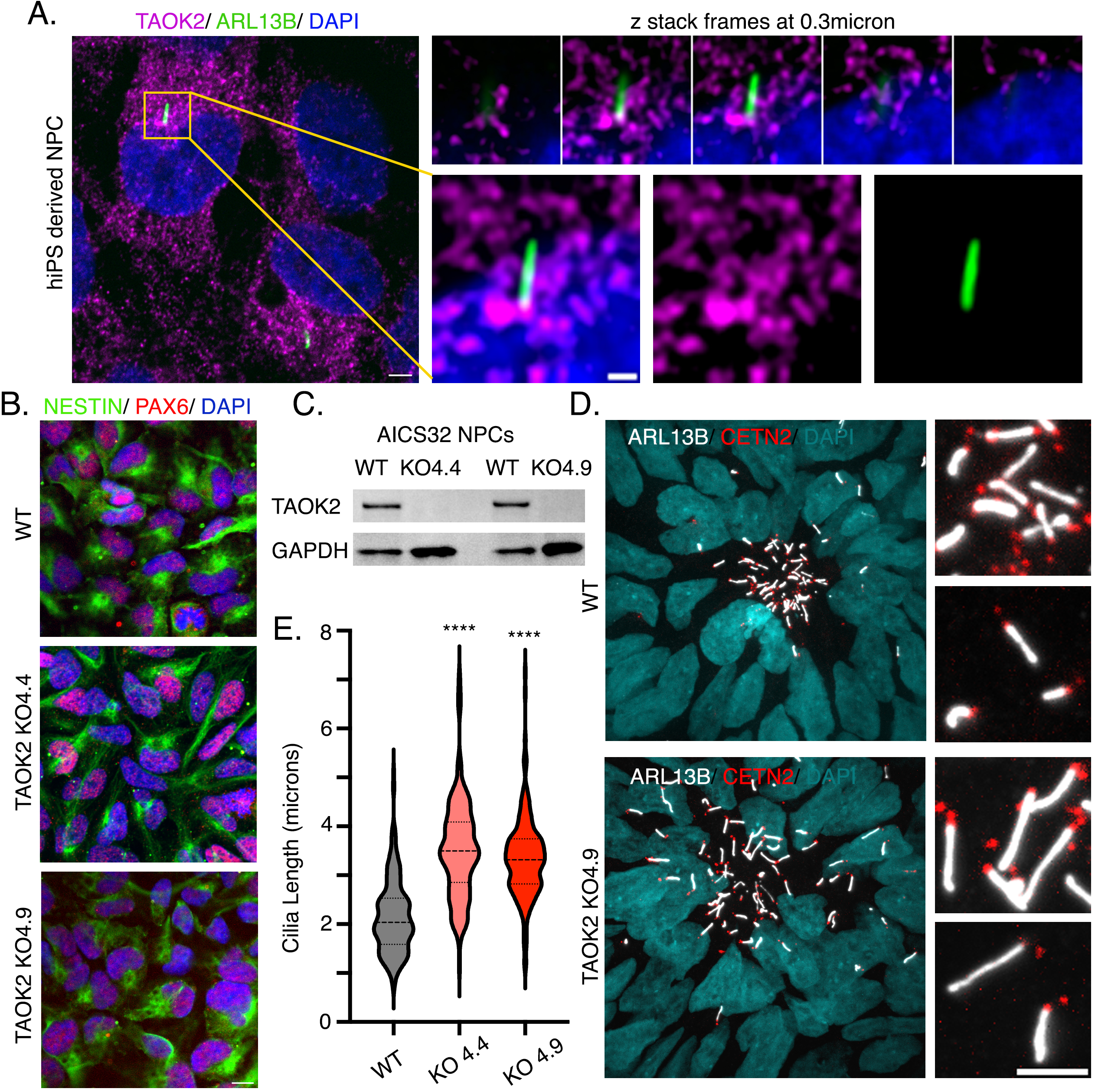
TAOK2 kinase localizes to the base of the primary cilium and restricts cilium elongation **(A)** Human iPS derived NPC stained for endogenous TAOK2 (magenta) and cilia marker ARL13B (green) shows ER localization of TAOK2 as well as its localization at the base of the primary cilia. Nuclei are stained with DAPI. Scale bar is 5μm. Magnified inset is shown on the right with individual frames of the z-stack acquired at 0.3μm are shown. Scale bar is 1μm. **(B)** Two distinct TAOK2 knockout iPS clones 4.4 and 4.9 were differentiated into NPCs and immunostained for dorsal progenitor markers PAX6 and NESTIN. Scale bar is 5μm. **(C)** Western blot analyses of protein abundance from TAOK2 KO AOK1 **(D)** Neural rosettes from control and TAOK2 KO cells expressing RFP-Centrin2 (red) were stained for ARL13B (white). Nuclei are shown in cyan (DAPI). Scale bar is 3μm. **(E)** Quantification of cilium length in control as well as TAOK2 KO 4.4 and KO 4.9 NPCs immunostained with ARL13B.

### TAOK2 Knockout Neural Progenitors Exhibit Ciliary Trafficking Deficits

Dedicated trafficking machinery comprised of the Intraflagellar Transport (IFT) proteins that mediate anterograde and retrograde motor-based transport of cargo in and out of the primary cilium^37,38^. We first tested whether the centrosomal scaffold protein Pericentrin, which is important for trafficking of IFT bound cargo^39,40^ and for cilium elongation^41^, was disrupted in TAOK2 KO NPCs. WT and TAOK2 KO NPCs were fixed and immunostained for endogenous Pericentrin (PCNT) and ciliary membrane marker ARL13B (Figure 5A). Mean fluorescence intensity of Pericentrin within a 10µm radius of the CETN2-RFP tagged centrosomes was significantly increased in TAOK2 KO4.4 (mean=336, n=220, S.E.M=23.22) and TAOK2 KO4.9 NPCs (mean=354.4, n=220, S.E.M=29.60) compared to WT NPCs (mean=204, n=220, S.E.M=8.10) (Figure 5B). Next, we tested whether intraflagellar transport is maintained in TAOK2 KO cells. We immunostained WT and TAOK2 KO NPCs with the anterograde IFT-B component IFT88, to measure kinesin-mediated trafficking to the cilia tip. WT and TAOK2 KO NPCs were fixed and immunostained for endogenous IFT88 (yellow) and ciliary membrane marker ARL13B (magenta) (Figure 5C). IFT88 particles are present at the base of the cilia and distributed along the length of the cilia, indicating normal transport. Accumulation at the cilia tip typically occurs in IFT-A mutants and revealing defective balance between retrograde and anterograde transport^42^. In WT NPCs, 9.76% and 16.36% of cells had IFT88 enriched at the ciliary base and tip, respectively, while in a majority (73.86%) of cells, IFT88 was uniformly distributed along the ciliary length (n=482). In contrast, IFT88 was enriched in the cilia tip in 61.95% of TAOK2 KO4.4 and 52.8% of TAOK2 KO4.9 NPCs, while it was uniformly distributed in 30.61% of TAOK2 KO4.4 and 35.42% of TAOK2 KO4.9 NPCs (n=460 and 595, respectively) (Figure 5D). These data show that TAOK2 regulates Pericentrin and IFT88 trafficking in the primary cilium, in absence of which there is increased centrosomal Pericentrin and IFT88 accumulation at the distal primary cilium tip.

**Figure 5.**
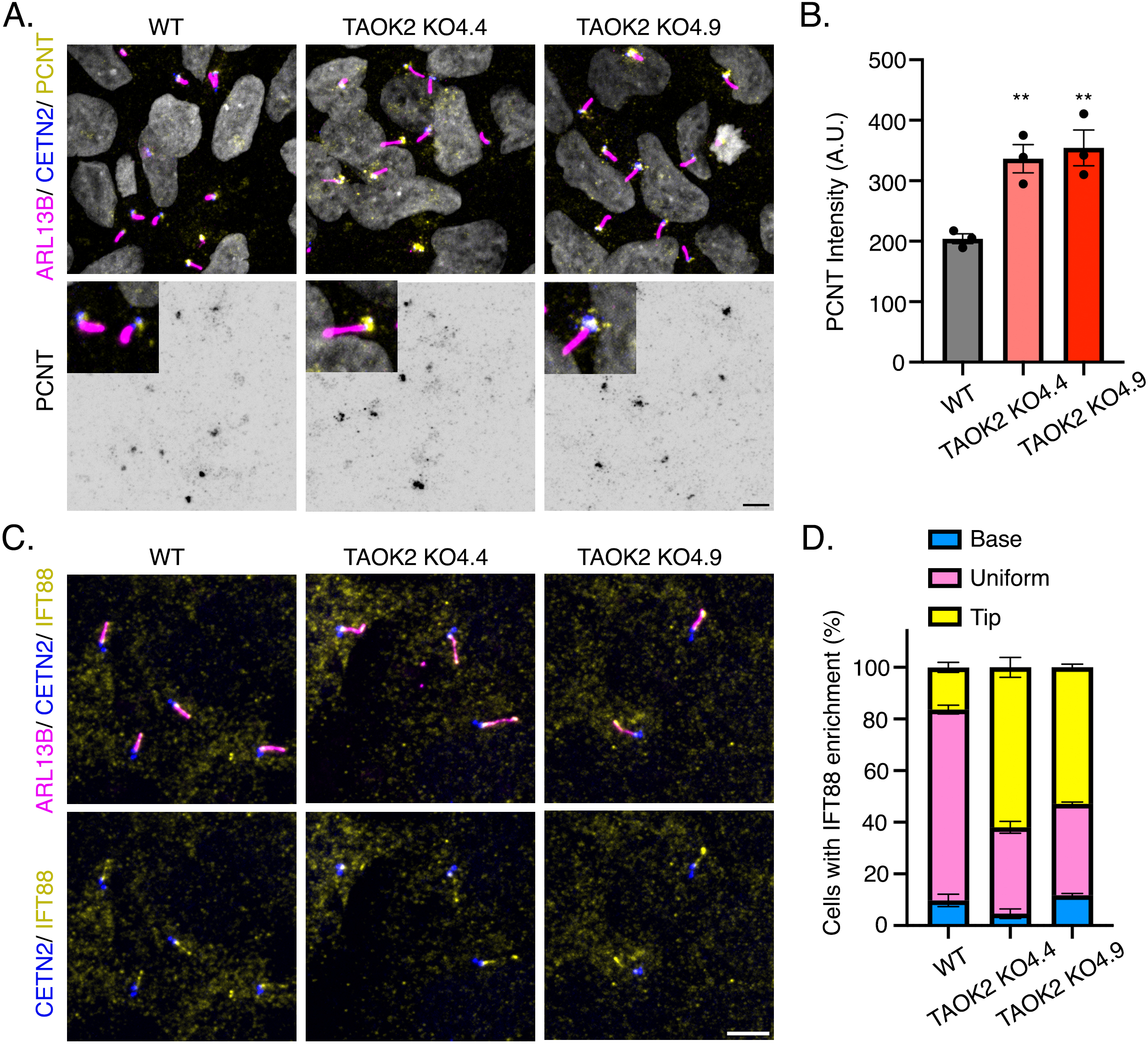
TAOK2 KO neural progenitors have increased centrosomal pericentrin and IFT88 accumulation at ciliary tip **(A)** NPCs derived from control and TAOK2 knockout 4.4 and 4.9 iPS cells endogenously tagged with Centrin 2 (blue) were immunostained for Pericentrin (PCNT) (yellow) and cilia marker ARL13B (magenta). Bottom row shows percentrin immunostaining in inverted grayscale. Scale bar is 5μm. **(B)** Fluorescence intensity of PCNT staining within a 10micron radius around the centrosome marked by centrin2 (blue) was measured in control and TAOK2 knockout NPCs. **(C)** NPCs derived from control and TAOK2 knockout 4.4 and 4.9 iPS cells endogenously tagged with Centrin 2 (blue) were immunostained for IFT88 (yellow) and cilia marker ARL13B (magenta). Bottom row shows merge image of IFT88 (yellow) with endogenous Centrin-2 (blue) that marks the base of the primary cilium. Scale bar is 5μm. **(D)** Percent of cilia with IFT88 enrichment at the cilium base, cilium tip and percent cilia with uniform distribution of IFT88 along the length of the cilia is plotted.

### TAOK2 Catalytic Activity is Required for its Role in Cilium Length Control

TAOK2 is a catalytically active kinase belonging to the serine threonine STE20 family^26^. The kinase activity of TAOK2 is cell cycle regulated, with a dramatic increase in activity during mitotic phase^31^, a cell cycle phase where the cilium is progressively reabsorbed. To determine whether TAOK2 mediates cilium length control through phosphorylation, we tested whether kinase activity of TAOK2 was required for its role. WT NPCs with endogenously tagged Centrin-2 were treated with control vehicle DMSO or the small molecule TAO kinase inhibitor CP43 at 10μM for 3hr. CP43 is a well characterized TAOK2 inhibitor with an IC50 =15nM^32,43^. After drug treatment, cells were fixed and immunostained for cilia marker ARL13B. We found that NPCs treated with CP43 showed a robust increase in cilia length (mean= 3.13µm, n=100, S.E.M.= 0.06) compared to DMSO treated controls (mean= 2.29µm, n=100, S.E.M.=0.07) indicating that TAOK2 kinase activity is important for its role in cilia length control (Figure 6A and 6B). These data reveal that TAOK2 catalytic activity controls cilium length, and its acute inhibition leads to increased cilia length, phenocopying the effect of TAOK2 knockout.

**Figure 6.**
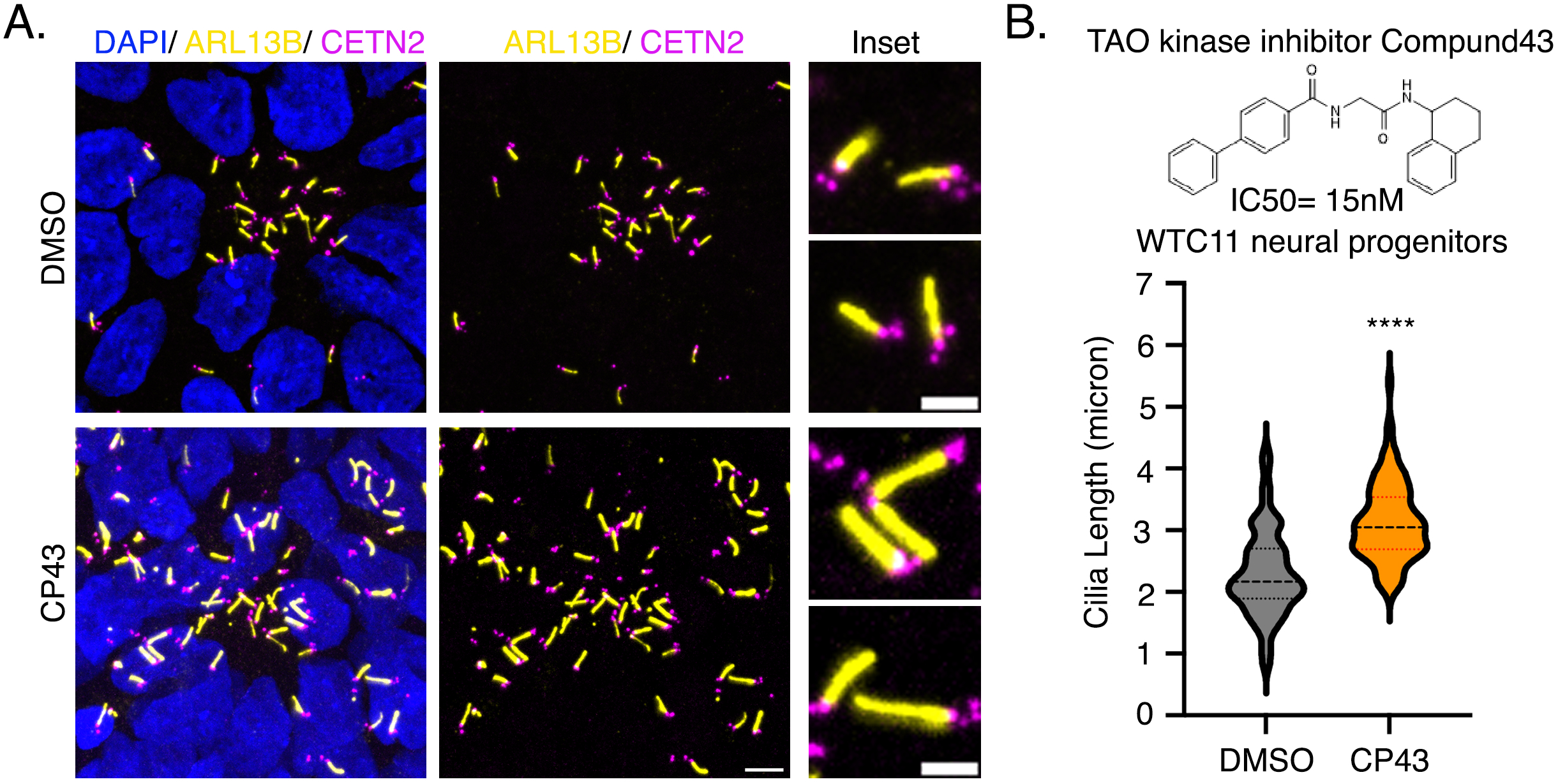
TAOK2 catalytic activity is important for its role in ciliary growth restriction **(A**) NPCs derived from control WTC11 iPSC (AICS0032) endogenously tagged with Centrin 2 (magenta) were treated with DMSO or TAO kinase inhibitor CP43 (1μM for 3 hr), fixed and immunostained for ARL13B (yellow) and nuclear dye DAPI (blue). Scale bar is 5μm. Inset on right shows magnified representative cilia treated with DMSO or CP43. Scale bar is 2μm. (B) Schematic shows the chemical structure and IC50 of CP43. Cilia length in NPCs treated with vehicle DMSO or CP43 is plotted.

## DISCUSSION

Ciliopathies, human diseases caused by mutations in genes affecting centrosomes and primary cilia, are commonly associated with severely impacted brain development^21,22^. Further, disruption in cilia length has been observed in several neurological disorders. Patients with Down syndrome, for example, can exhibit shorter cilia, associated with aberrant centrosomal accumulation of Pericentrin that leads to impaired trafficking to the cilium^44^. Aberrant ciliogenesis is also seen in patients with focal malformations of cortical development (FMCD) caused by somatic activating mutations in mTOR with developmental delay, epilepsy, and autism^45^. Seckel Syndrome, caused by mutations in the centrosomal-P4.1-associated protein (CPAP) is associated with microcephaly. Seckel Syndrome patient-derived NPCs exhibit extremely long cilia length and undergo premature neuronal differentiation^46^. At the outset, several lines of evidence pointed to a potential role of ciliary dysfunction in 16p11.2 CNVs. First, transcriptomic profiles of individuals carrying reciprocal 16p11.2 CNVs were enriched for genes associated with ciliopathies^47^. Second, radial glial cells in cerebral organoids generated from 16p11.2 carriers show differentially expressed cilium assembly genes^15^, and lastly, 16p11.2 CNV patients exhibit BMI and metabolic imbalances^1,6,7,48^, phenotypes often found in ciliopathy patients^24,25,49^. In this study, our unbiased proteomic analyses of human iPSC-derived neural progenitors from 16p11.2 deletion and duplication carriers revealed that altered phosphoproteins in 16p11.2 CNVs were enriched for proteins with a centrosomal and cilia related function. Reciprocal changes in cilia length in NPCs derived from 16p11.2 deletion and duplication carriers were identified in our study. Whether reciprocal cilia length deficits underlie the opposing changes in brain size and BMI in 16p11.2 CNV carriers is an important future direction of investigation. Additionally, given the incomplete penetrance of 16p11.2 clinical symptoms^2,12^, if such an association of cilia length is made with brain size or BMI, cilium length alteration, could serve as a valuable biomarker for 16p11.2 CNV carriers.

During development of the mammalian neocortex, neuroepithelial cells of the neural tube dramatically expand and differentiate into radial glial cells^22,50^. These polarized radial glial cells are organized with their apical plasma membranes forming the luminal side of the neural tube, such that the primary cilia extend into the ventricular cerebrospinal fluid rich in signaling molecules^51^. Enriched within the primary cilium are numerous signaling receptors that sense growth factors and ligands important for brain development such as insulin-like growth factors, bone morphogenic proteins, sonic hedgehog, retinoic acid and lysophosphatic acid^22,24^. Cilia are dynamic and undergo length extension, retraction and tip-excision during the cell cycle. Cilia length dynamics are observed during neuronal differentiation, with the circadian rhythm and in response to growth factors through mechanisms that are not well defined^37,38,41,52–56^. Through its critical signaling functions and dynamic regulation of length that dictates receptor availability, the primary cilium serves as key regulator of neural stem cell proliferation, neuronal differentiation, migration as well as site of specialized axon-cilium synapse^22,57–59^. Whether and how any of these cilium dependent processes are impacted in 16p11.2 CNVs remains to be investigated.

An important function of the primary cilium is energy homeostasis, and mutations in human genes that perturb ciliary function cause obesity and metabolic defects^25^. Melanocortin 4 receptor (MC4R) colocalizes with adenylyl cyclase III at the primary cilium in a subset of neurons in the hypothalamus. MC4R mutations impair ciliary localization and inhibition of adenylyl cyclase signaling at the primary cilia of hypothalamic neurons, leading to increased body weight^60^. Conditional knockout of leptin receptor in neural stem cells leads to obesity in mice, leptin-induced cilia assembly is essential for sensing satiety signals by hypothalamic neurons^55^. The genetic basis of obesity in 16p11.2 deletion carriers is an area of active investigation. Our results demonstrating reciprocal changes in cilia length in deletion and duplication carriers with opposing BMI changes brings the intriguing possibility that aberrant ciliary signaling in 16p11.2 CNV underlie the changes in bioenergetics in these patients. Importantly, phenome-wide association studies between imputed expressions of individual 16p11.2 genes and over 1500 health traits found genes TMEM219, SPN, TAOK2, INO80E within 16p11.2 associated with BMI and obesity^61^. Our results show that TAOK2 gene dosage regulates cilia length and that neural progenitors lacking TAOK2 have elongated cilia and increased centrosomal Pericentrin and accumulation of IFT88 at distal cilia tip. Notably, TAOK2 is the only gene within the 16p11.2 locus, single nucleotide missense mutations in which have been associated with obesity^62,63^. Whether and how the ciliary role of TAOK2 is associated with its contribution of metabolic dysfunction is an exciting direction for future investigation.

### Limitations of the Study

Our study identified opposing ciliary length defects in neural progenitors derived from three distinct carriers of 16p11.2 deletion and duplication, and determined the genetic basis for these alterations. Whether these ciliary deficits in exist broadly in other cell types within the brain, and in other tissues needs to be examined. While our study provides compelling evidence for the dosage dependent modulation of cilia length by TAOK2, and that phosphorylation by TAOK2 mediates cilia length control, precise mechanisms remain to be elucidated. Chemical genetics approaches to identify direct phosphorylation targets of TAOK2 revealed several proteins that play important roles in cilia structure and function, including Septin 7, HDAC6, MST3, CEP170 and ASAP1^27^. TAOK2, through its C-terminal tail interacts with microtubules and EB1 proteins^31^, both of which have important structural roles in cilia elongation. How these distinct TAOK2 dependent pathways uniquely or in concert determine cilia length is an important area of inquiry. Finally, in light of the inherent pleiotropy of TAOK2 conferred by several functional domains, it will be critical to dissect key molecular elements that determine its ER-membrane, centrosomal, and ciliary functions.

## METHODS

### Induced pluripotent stem cell culture and neuronal differentiation

All iPSC lines (16p11.2 control, duplication, and deletion; WTC11, AICS0032 WT and TAOK2 KO4.4 and TAOK2 KO4.9) were differentiated into NPCs as reported previously^14^. Stem cells were treated with 1U/mL Dispase, lifted using a cell scraper, resuspended in mTeSR media (STEMCELL Technologies) containing 10μm ROCK Inhibitor (Selleck Chemical), and then transferred to uncoated T25 flasks for embryoid body (EB) formation. After 24-48 hours (Day 0), EBs were gravity settled and transferred to new uncoated T25 flasks in neural media (DMEM/F12 (Gibco), 1% N-2 supplement (Gibco), 1% Non-essential amino acids (Gibco), 2μg/mL Heparin, 1% Penicillin/Streptomycin [Gibco]) containing small molecule TGF-β/SMAD inhibitors, SB431542 (5μM; Peprotech) and LDN193189 (0.25μM; Peprotech). On Day 3, EBs were transferred to Matrigel (1:50 dilution; Corning)-coated 6 well plates in fresh neural media with SB and LDN. Media was changed with neural media without SB and LDN every other day or as needed until Day 11 for formation of neural rosettes. On Day 11, neuroepithelia were mechanically lifted using cell scraper and transferred to uncoated T25 flasks in neural media. Neurospheres were fed with neural media every other day, or as needed, until Day 25. On Day 25, neurospheres were plated onto Matrigel-coated plates in STEMDiff™ Neural Progenitor Media (STEMCELL Technologies) and NPCs were obtained.

### Proteomic sample preparation

Aliquots of patient derived neural progenitor cells were digested with trypsin for proteomic analysis as follows. Confluent 75mL flasks of NPCs were washed with PBS and then homogenized in 1ml 8M Urea, using a probe sonicator (ThermoFisher Scientific). Total protein content in the extracts was determined using a bicinchoninic acid protein assay kit (Micro BCA Protein Assay Kit, Thermo Scientific). Aliquots containing 500μg of protein in 590μl 8M urea were added with 236μl of 200 mM ammonium bicarbonate buffer and phosphatase inhibitors (23μl of each Sigma Phosphatase Inhibitor Cocktails 1 and 3), then treated with 5mM DTT at 56°C for 15 minutes, followed by a 30-minute incubation at room temperature in the dark with 7.5 mM iodoacetamide. For tryptic digestion, the samples were then diluted 4-fold with 100 mM ammonium bicarbonate to reduce urea concentration to 2 M and then added 5% (W/W) modified trypsin (Promega, Madison, WI). The pH was adjusted to 8.0 with 250 mM ammonium bicarbonate, and the samples were incubated 12 h at 37 °C. After that, another aliquot of trypsin was added (2% W/W) and digested for additional 6 hours. After this, samples were acidified to a final concentration of 5% formic acid. The digests were then desalted using a MAX-RP Sep Pak ® classic C18 cartridge (Waters) following the manufacturer’s protocol. Sep Pak eluates were dried-evaporated in preparation for labeling with TMT-10 plex reagents.

### TMT Labelling

Dried samples were labeled according to TMT-10 label plex kit instructions (ThermoFisher Scientific), with some modifications. Briefly, samples were resuspended in 100μl 0.1M triethylammonium bicarbonate pH 8.0. TMT reagents (600μg) were dissolved in 41μl acetonitrile, and added to the samples. After incubation for 1 h at room temperature samples were quenched with 8µl 5% hydroxylamine, then all 10 samples combined and partially evaporated in speedvac until volume was around 5µl. 100ul 1% formic acid was added, and then peptides desalted using a C18 SepPak. The Sep Pak eluate was dried in preparation for phosphopeptide enrichment.

### Phosphopeptide enrichment

Phosphopeptide enrichment was performed in an AKTA Purifier (GE Healthcare, Piscataway, NJ) using 5µm titanium dioxide (TiO2) beads (GL Sciences, Tokyo, Japan) in-house packed into a 2.0mm x 2 cm analytical guard column (Upchurch Scientific, Oak Harbor, WA). Combined TMT labelled tryptic digests were resuspended in 240μl buffer containing 35% MeCN, 200 mM NaCl, 0.4% TFA and loaded onto the TiO2 column at a flow rate of 2 ml/min. The column was then washed for 2 min with 35% MeCN, 200 mM NaCl, 0.4% TFA to remove non phosphorylated peptides. Phosphopeptides were eluted from the column using 1M Potassium Phosphate Monobasic (KH2PO4) at a flow rate of 0.5 ml/min for 30 min directly onto an on-line coupled C18 macrotrap peptide column (Michrom Bioresources, Auburn, CA). This column was washed with 5% MeCN, 0.1% TFA for 14 min and the adsorbed material was eluted in 400 µl of 50% MeCN, 0.1% TFA at a flow rate of 0.25 ml/min. The eluate was solvent evaporated and then resuspended in 240 ml 20 mM ammonium formiate pH 10.4 for fractionation of the peptide mixture by high pH RP chromatography.

### High pH Reverse Phase Chromatography

The phosphopeptide enriched sample (Solvent evaporated TiO2 chromatography eluate) was resuspended in 240μl of 20mM ammonium formate pH 10.4 for fractionation of the peptide mixture by high pH RP chromatography using a Phenomenex Gemini 5u C18 110A 150 x 4.60 mm column, operating at a flow rate of 0.550 mL/min. Buffer A consisted of 20 mM ammonium formate (pH 10), and buffer B consisted of 20 mM ammonium formate in 90% acetonitrile (pH 10). Gradient details were as follows: 1 % to 30% B in 49 min, 30% B to 70% B in 4 min, 70% B down to 1% B in 4 min. Peptide-containing fractions were collected, combined into 25 final fractions for LC-MS/MS analysis, evaporated, and resuspended in 20μl 0.1% formic acid for mass spectrometry analysis.

### Mass Spectrometry Analysis

Peptide resuspended in 0.1% formic acid were injected (5μl) onto a 2µm x 75 µm x 50 cm PepMap RSLC C18 EasySpray column (Thermo Scientific). For peptide elution, 3-hour water/acetonitrile gradients (2–25% in 0.1% formic acid) were used, at a flow rate of 200 nl/min. Samples were analyzed in an QExactive Plus Orbitrap (Thermo Scientific) in positive ion mode. MS spectra were acquired between 350 and 1500 m/z with a resolution of 70000. For each MS spectrum, the 10 higher intensity multiply charged ions over the selected threshold (1.7E4) were selected for MS/MS (apex trigger 1 to 10s) with an isolation window of 1.0 m/z. Precursor ions were fragmented by HCD using stepped relative collision energies of 25, 35 and 40 to ensure efficient generation of sequence ions as well as TMT reporter ions. MS/MS spectra were acquired in centroid mode with resolution 70000 from m/z=100. A dynamic exclusion window was applied which prevented the same m/z from being selected for 10s after its acquisition.

### Peptide and protein identification and quantitation

Peak lists were generated using PAVA in-house software^34^. All generated peak lists were searched against the human subset of the SwissProt database (SwissProt.2016.9.6, 20198 entries searched), using Protein Prospector^33^ with the following parameters: Enzyme specificity was set as Trypsin, and up to 2 missed cleavages per peptide were allowed. Carbamidomethylation of cysteine residues, and, in the case of TMT labelled samples, TMT10plex labeling of lysine residues and N-terminus of the protein, were allowed as fixed modifications. N-acetylation of the N-terminus of the protein, loss of protein N-terminal methionine, pyroglutamate formation from of peptide N-terminal glutamines, oxidation of methionine, were allowed as variable modifications. Mass tolerance was 20 ppm in MS and 30 ppm in MS/MS. The false positive rate was estimated by searching the data using a concatenated database which contains the original SwissProt database, as well as a version of each original entry where the sequence has been randomized. A 1% FDR was permitted at the protein and peptide level. For quantitation only unique peptides were considered; peptides common to several proteins were not used for quantitative analysis. For TMT based quantitation, relative quantization of peptide abundance was performed via calculation of the intensity of reporter ions corresponding to the different TMT labels, present in MS/MS spectra. Intensities were determined by Protein Prospector. Summed intensities of the reporter ions (each TMT channel) for all peptide spectral matches (PSMs) were used to normalize individual (sample specific) intensity values. For each PSM, relative abundances were calculated as ratios vs the average intensity levels in the channels corresponding to control (normal 16p11.2 individuals) samples. PSMS ratios were aggregated to peptide level using median values of the log2 ratios. Statistical significance was calculated with a 2-tailed t-test.

### NPC transfection and immunocytochemistry

NPCs grown on coverslips were washed with phosphate buffer saline (PBS) once, fixed with 4% paraformaldehyde (PFA), and 4% sucrose solution for 20 minutes at room temperature, and then washed thrice with PBS. Fixed NPCs were then incubated with blocking buffer (10% Normal Goat Serum, 200mM Glycine pH 7.4, 0.25% Triton X-100 in PBS) for one hour at room temperature. Cells were incubated with primary antibodies in blocking buffer overnight at 4℃, followed by six 5-minute PBS washes and incubation with secondary antibodies at 1:1000 dilution in blocking buffer overnight at 4℃. DAPI was included in the secondary antibody mixture at 1:2000 dilution. Coverslips were washed six to ten times with PBS and mounted on slides with Fluoromount-G (Electron Microscopy Sciences).

For transfection with expression constructs and shRNA plasmids, NPCs were transfected using Lipofectamine™ Stem Transfection Reagent (Invitrogen) according to the manufacturer’s protocol.

### DNA extraction, purification, and sequencing

Genomic DNA was extracted from iPSCs and NPCs using QuickExtract™ DNA Extraction Solution (Lucigen) according to the manufacturer’s protocol. Regions of interest were PCR amplified with the appropriate primers and purified using ExoSAP-IT™ Express PCR Product Cleanup Reagent (Applied Biosystems) according to the manufacturer’s protocol. Purified PCR products were sent to GeneWiz for sequencing with the appropriate primers.

### Protein extraction and western blot

NPCs or iPSCs were lysed with HKT buffer (25mM HEPES pH 7.2, 150mM KCl, 1% Triton X-100, 2mM DTT, 1x protease inhibitor (Roche, cOmplete™ EDTA free), 1mM EDTA) and collected using a cell scraper, followed by addition of Pierce™ Universal Nuclease for Cell Lysis (Thermo Scientific) and lysed with several passages through a sterile 25-gauge syringe needle. NuPAGE™ LDS Sample Buffer (Invitrogen) containing 125mM DTT was added to the sample which was then heated for 10 minutes at 95℃. Samples were run on NuPAGE™ 4-12% Bis-Tris Polyacrylamide gels (Invitrogen) with NuPAGE™ MOPS running buffer (Invitrogen) at 165V for 20 minutes and then 175V for 50 minutes. Gels were transferred to Immobilon-P membrane (Millipore) at 100V for 1 hour. Blots were blocked with 5% BSA blocking buffer and probed with indicated antibodies overnight at 4℃ followed by 3hr incubation with HRP conjugated secondary antibodies at 1:5000 dilution at room temperature. The blots were imaged with Pierce™ ECL Western Blotting Substrate (Thermo Scientific) using ChemiDoc Imager (Bio-Rad).

### Generation of TAOK2 knockout stem cell line

Two independent TAOK2 knockout cell lines were generated using CRISPR/Cas9 genome editing in a AICS0032 stem cell line. Two guides were designed using Synthego guide design tool (https://www.synthego.com) to target coding exon 2 and 4. Guides were cloned into plasmid CrisprV2pSpCas9(BB)-2A-Puro (PX459) V2.0 (Addgene Plasmid #62988) with Puromycin resistance. Cells were passaged in single cell suspension and plated at 50% confluence. Cultures were then transfected with lipofectamine 2000 reagent (Invitrogen) and 2mg of each of the 2 guides used per KO line. Cells were then selected with Puromycin for 48hr to select for transfected cells. Edited cells were expanded into single cell colonies. Genomic DNA was extracted and the region around the cutting site was PCR amplified to send for sequencing. Successful TAOK2 knockout clones 4.4 and 4.9 were confirmed by Sanger sequencing analysis, and absence of encoded protein was validated using western blot. TAOK2 KO lines 4.4 and 4.9 were confirmed by karyotyping for absence of large chromosomal alterations.

### Confocal Microscopy

Imaging was performed on a Nikon Ti2 Eclipse-CSU-X1 confocal spinning disk microscope equipped with four laser lines (405nm, 488nm, 561nm, 670nm) and an sCMOS Andor camera for image acquisition. The microscope was caged within the OkoLab environmental control setup enabling temperature and CO2 control during live imaging. Imaging was performed using Nikon 1.49 100x Apo 60X or 40X oil objective lenses. All image analyses and quantification of fluorescence intensity was performed using Fiji ImageJ2 version2.3.

### Statistics

All statistics were performed in GraphPad software Prism10.0. Multiple groups were analyzed using ANOVA, while two group comparisons were made using unpaired t test unless otherwise stated. Statistically *p* value less that 0.05 was considered significant. All experiments were done in triplicate, and experimental sample size and p values are indicated with the corresponding figures.

## Supporting information

combined supplementary

## Acknowledgments

We are grateful for funding provided by the National Institutes of Mental Health (R01MH121674) to S.Y. and the Jaconette Tietze Award to S.Y. We extend our gratitude to the patients and families of 16p11.2 CNV carriers who have shared their skin samples to support research, and funding from SFARI(345471) to L.A.W for generation of the iPSC lines. Mass Spectrometry was provided by the Mass Spectrometry Resource at UCSF (A.L.B, Director) supported by the Dr. Miriam and Sheldon G. Adelson Medical Research Foundation and the UCSF Program for Breakthrough Biomedical Research.

## Author contributions

A.F. designed and performed shRNA and overexpression screening experiments and generated the TAOK2 knockout iPSC lines. S.B. characterized the TAOK2 KO iPSC lines, differentiated them into neuronal lineage and performed all cilia characterization experiments on the KO lines. Cilia database analyses of proteomic data and molecular biology experiments were performed by M.C. Proteomics experiments were performed and analyzed by J.O.P under the supervision of A.L.B. Neuronal differentiation of 16p11.2 deletion and duplication as well as control iPS lines were performed by A.D. under the supervision of L.A.W. The study was designed and written by S.Y. and edited by all authors. All aspects of work, with exception of the proteomics, were supervised by S.Y.

## Declaration of interests

The authors declare no competing interests.

## Notes

### Competing Interest Statement

The authors have declared no competing interest.

