## Supplementary material for "TAOK2 Drives Opposing Cilia Length Deficits in 16p11.2 Deletion and Duplication Carriers": combined supplementary

### Supplementary Information

1. Supplementary Figures S1-S2
2. Supplementary Figure Legend
3. Supplementary Table Legend

### Supplementary Figure Legend

**Figure S1. Human neuronal model of 16p11.2 deletion and duplication** (related to main figure 1 and 2)

(A) Table shows iPSC lines used in the study. Three lines were derived from unaffected individuals,16p11.2 duplication and deletion carriers each, and their age, sex and source is shown. (B) Workflow and timeline for derivation of dorsal forebrain neural progenitors from iPSCs using dual SMAD inhibition by LDN and SB. (C) Representative confocal images of NPCs derived from control and 16p11.2 CNV carriers and immunostained with dorsal forebrain marker PAX6 (red) and co-stained with DNA dye DAPI (blue). Scale bar is 10μm.

**Figure S2. Generation of TAOK2 knockout human iPSC and NPCs using genome editing** (related to main figure 4)

(A) Schematic depicts the CETN2 genomic locus and position where mTagRFP was inserted using the indicated guideRNA to generate the AICS0032 stem cell line derived from WTC11. Data from Allen Institute. (B) Images show AICS0032 stem cells clone 19 expressing mTagRFP-Centrin 2. Centrioles, both mother and daughter are visible under the microscope. Data from Allen Institute website. (C) Expression plasmid backbone and site of guideRNA used to generate TAOK2 KO cell lines is shown. (D) Genomic sequencing of TAOK2 KO stem cells, show the sequence near exon4 that was targeted. (E) Western blot shows cell lysate from WT and TAOK2 KO clones 4.4 and 4.9 immunoblotted with antibodies against TAOK2 and GAPDH. (F) Karyotype analyses of WT and TAOK2 KO 4.9. (G) WT and TAOK2 KO 4.4 and 4.9 iPSCs immunostained for pluripotency markers NANOG (magenta), OCT4(cyan) and nuclear dye DAPI (blue). Scale bar is 20μm.

### Supplementary Table Legend

**Table S1. TMT10Plex Proteomics to Discover Altered Phosphoproteins in 16p11.2 CNV carrier derived neural progenitor cells** (related to Figure 1)

List of TMT10plex reagent labelled phosphopeptides identified by mass spectrometry from 16p11.2 deletion, duplication and control iPSC derived NPCs.

**Table S2. Centrosomal and Ciliary Proteins are differentially phosphorylated in 16p11.2 CNV and control human NPCs** (related to Figure 1)

List of proteins aberrantly phosphorylated in 16p11.2 CNV iPSC derived NPCs queried against their presence of absence in the CCDB database and GeneCards Cilia search.

A.

| Line Name | Genotype | ASD Diagnosis | Sex | Age (yr) | Source |
|---|---|---|---|---|---|
| 0162D | Control | No | Female | 15 | Published, Weiss Lab |
| 0165D | Control | No | Female | 12 | Published, Weiss Lab |
| 0119B | Control | No | Male | 44 | Published, Weiss Lab |
| 03C | 16p11.2 Duplication | Yes | Male | 5.7 | SFARI ID 14724.x5 |
| 04C | 16p11.2 Duplication | Yes | Male | 35.5 | SFARI ID 14723.x10 |
| 05C | 16p11.2 Duplication | Yes | Female | 2 | SFARI ID 14770.x7 |
| 06C | 16p11.2 Deletion | Yes | Female | 14 | SFARI ID 14753.x5 |
| 07C | 16p11.2 Deletion | Yes | Male | 8 | SFARI ID 14739.x3 |
| 08C | 16p11.2 Deletion | Yes | Female | 13 | SFARI ID 14755.x17 |

B.

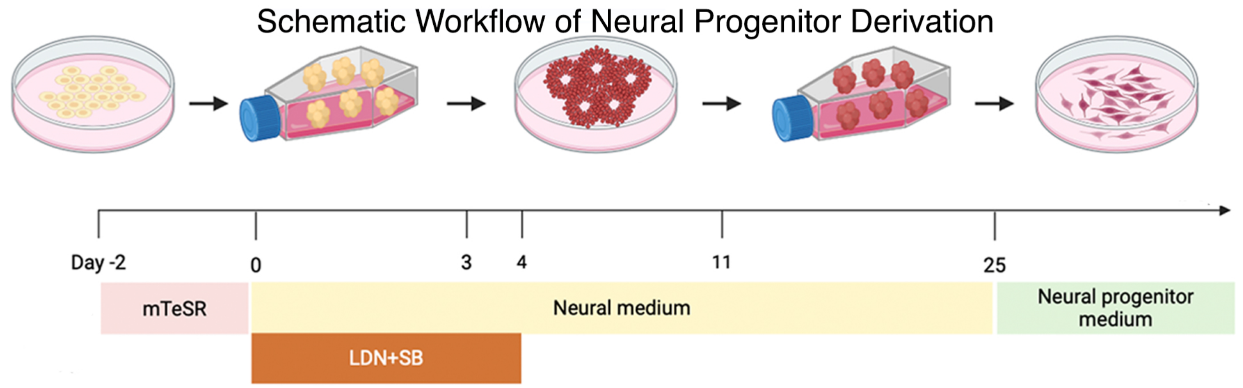


C.

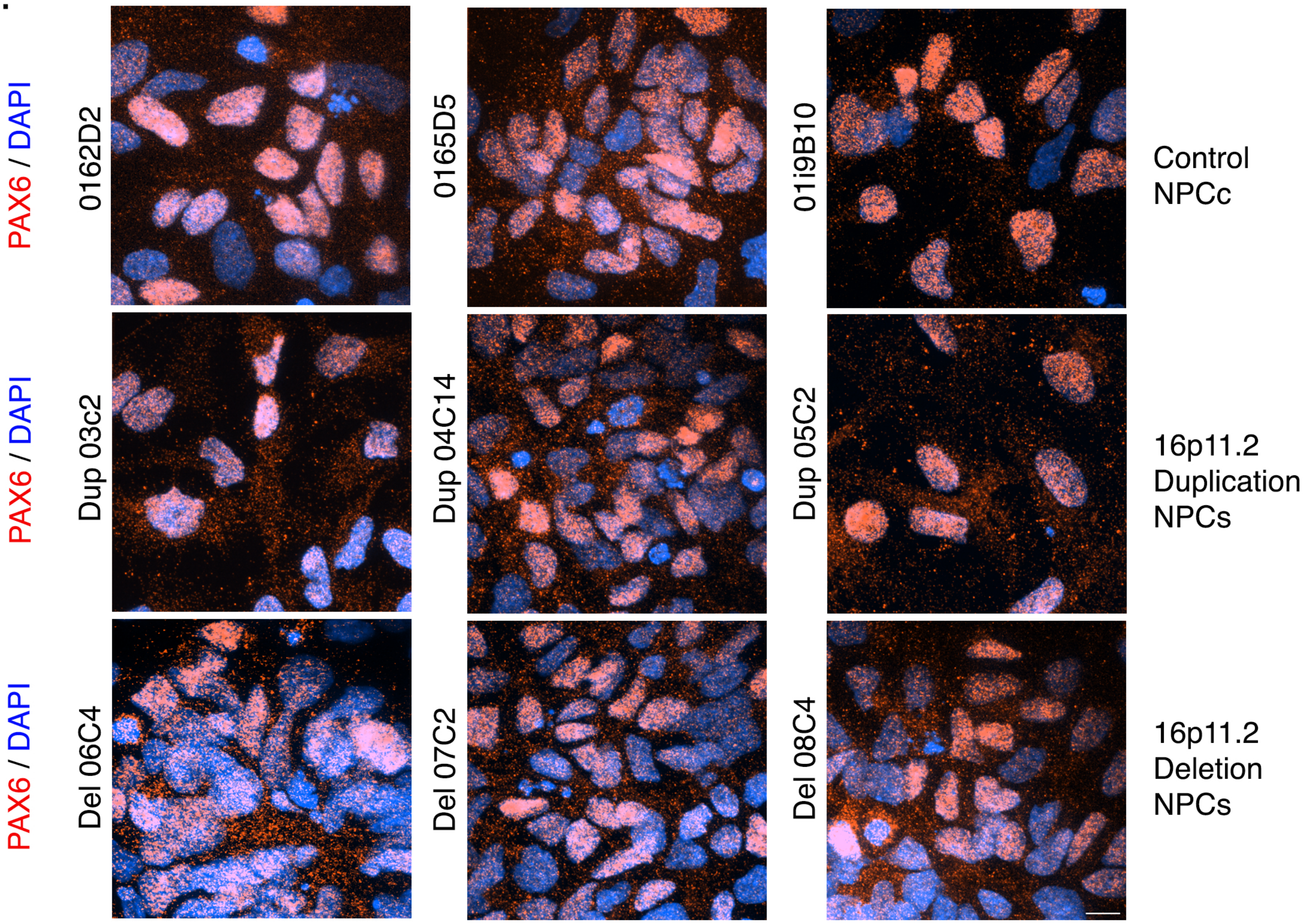


Figure S1

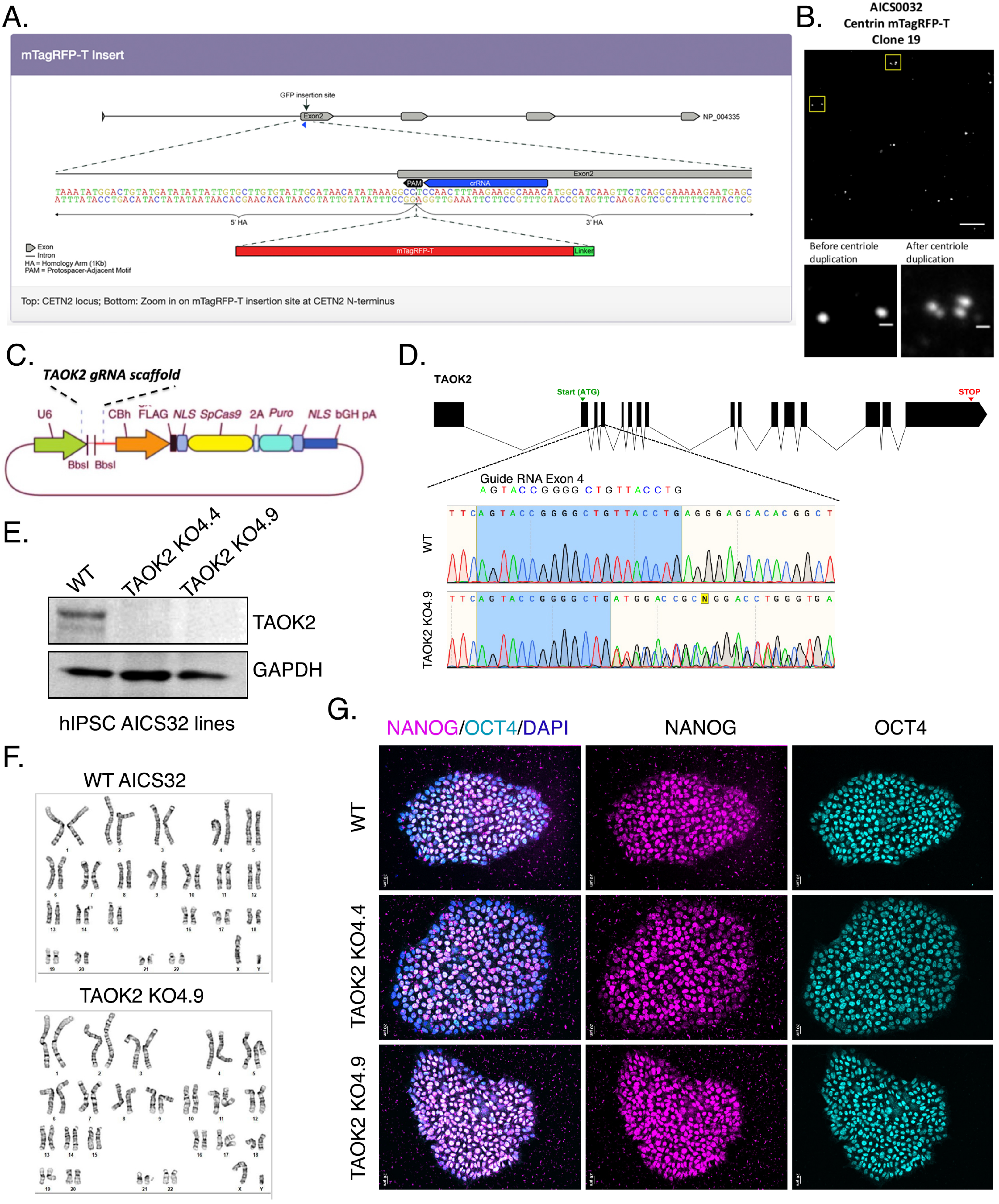

A.
mTagRFP-T Insert
GFP insertion site
Exon2
NP_004335
PAM
crRNA
5' HA
3' HA
mTagRFP-T
Linker
Exon
Intron
HA = Homology Arm (1Kb)
PAM = Protospacer-Adjacent Motif
Top: CETN2 locus; Bottom: Zoom in on mTagRFP-T insertion site at CETN2 N-terminus
B.
AICS0032
Centrin mTagRFP-T
Clone 19
Before centriole duplication
After centriole duplication
C.
TAOK2 gRNA scaffold
U6
CBh
FLAG
NLS
SpCas9
2A
Puro
NLS
bGH pA
BbsI
BbsI
D.
TAOK2
Start (ATG)
STOP
Guide RNA Exon 4
AGTACCGGGGCTGTTACCTG
WT
TAOK2 KO4.9
E.
WT
TAOK2 KO4.4
TAOK2 KO4.9
TAOK2
GAPDH
hIPSC AICS32 lines
F.
WT AICS32
TAOK2 KO4.9
G.
NANOG/OCT4/DAPI
NANOG
OCT4
WT
TAOK2 KO4.4
TAOK2 KO4.9


Figure S2
